# Acetyl-CoA availability regulates neuronal metabolism, growth, and synaptic activity

**DOI:** 10.64898/2026.01.20.700435

**Authors:** Eric R. McGregor, Cassandra J. McGill, Nicholas L. Arp, Josef P. Clark, Dominique A. Baldwin, Di Kuang, Gonzalo Fernandez-Fuente, Yun Hwa Choi, Keenan S. Pearson, Jason K. Nagorski, Judith A. Simcox, Jing Fan, Luigi Puglielli, Rozalyn M. Anderson

## Abstract

The metabolite acetyl-CoA plays a central role in cellular metabolic homeostasis. As part of the secretory pathway, acetyl-CoA is imported into the endoplasmic reticulum (ER) by a membrane-bound transporter AT-1 (SLC33A1). AT-1 has been linked to peripheral neuropathy (heterozygous mutations), developmental delay with premature death (homozygous mutations) and intellectual disability with progeria (duplication). These phenotypes can be reproduced in the mouse. Here, we show that AT-1 overexpression in primary neurons impacts diverse phenotypes related to neuronal function and plasticity. At the gene level, AT-1 induces brain aging signatures, and key differences in ribosomal and synaptic processes were identified in both the transcriptome and the proteome. Changes in mitochondria-associated pathways were reflected in an increase in expression of mitochondrial master regulator PGC-1α and its target genes. Functionally, marked differences in mitochondrial membrane potential, architecture, and respiration were detected. Tracing experiments indicated altered glucose utilization in glycogen storage and nucleotide production. Shifts in redox metabolism were linked to differences in levels of NAD-dependent SIRT1 and CtBP2, with consequences for acetylated lysine modification. Depletion of lipid stores was associated with greater plasticity in fuel substrate utilization and a major shift in cellular lipid composition. These broad-scale changes in metabolism were coincident with reduced expression of synaptic proteins and reduced activity among synaptic networks, indicating that neuronal electrophysiology and network communication are coordinated at least in part through neuronal acetyl-CoA metabolism.

## Introduction

Neurons require efficient energy production to fuel synaptic communication. They also require efficient protein quality control mechanisms to deliver properly folded proteins to the synapse and other parts of the cell membranes. Neuronal function relies on coordinated regulation of metabolic, structural, and communication pathways, matching energy derivation to biosynthetic needs. It is not surprising then that neurodegeneration and autism spectrum disorder are associated with alterations in metabolism (Grimm & Eckert 2017; Khaliulin et al. 2025). We recently described overt changes in metabolism in the aging brain that were coincident with signatures of increased inflammation and immune dysfunction and with adverse changes in pathways associated with neuronal function (Souder et al. 2025). In neurons, metabolic activation impinges on proteostasis, growth, and processes linked to neurotransmission (McGregor et al. 2025). In cultured pre-adipocytes, even small changes in metabolism have profound effects on diverse cellular functions, from cell cycle to chromatin remodeling, from proteostasis to RNA processing (Miller, Clark, Martin, et al. 2019). Given the breadth of pathways and processes influenced by mitochondrial status, there is considerable motivation to go beyond the concept of “dysfunction” (Monzel et al. 2023) to advance understanding of the influence of metabolic status on neuronal function.

One of the critical molecules at the intersection of energetics and biosynthesis is acetyl-CoA. Acetyl-CoA is an ancient molecule that plays roles in metabolic sensing, regulation, and adaptation (Pietrocola et al. 2015). Acetyl-CoA is a product of glycolysis and lipid oxidation, a precursor of lipid biosynthesis, and a substrate for post-translational modification of diverse proteins, where acetylation influences function, chromatin accessibility, and protein stability. The dual identities of a metabolic intermediate and a functional modifier place acetyl-CoA at the pivot point between cellular catabolism and anabolism, influencing fuel preference through its impact on metabolic flux and energetic capacity as a post-translational modification and allosteric regulator.

In the brain, acetyl-CoA is similarly centrally poised: a role in maintaining equilibrium between anabolic and catabolic pathways within and among cell types has been proposed (Jankowska-Kulawy et al. 2022). Cytosolic acetyl-CoA is synthesized from citrate and CoA by ATP Citrate Lyase (ACLY) and from acetate and CoA by acetyl-CoA synthetase (ACSS2). It can then be shuttled by AT-1 (SLC33A1), an endoplasmic reticulum (ER) membrane-bound transporter (Jonas et al. 2010), into the ER lumen where it is used by two acetyltransferases for Nε-lysine acetylation of both cargo and resident proteins(Ko & Puglielli 2009). In mice, reduced cytosol-to-ER transport of acetyl-CoA via inactivation of AT-1 causes neurodegeneration of both central and peripheral nervous system (Peng et al. 2014), while increased cytosol-to-ER transport of acetyl-CoA causes behavioral abnormalities when AT-1 is overexpressed only in forebrain neurons (Hullinger et al., 2016) and a progeroid phenotype with reduced lifespan when AT-1 is overexpressed in the whole body (Peng et al. 2018).

A metabolic consequence of AT-1 manipulation is hinted at in the proteomics of brains from these mice, which show enrichment for mitochondrial proteins (Hullinger et al., 2016). Independent studies report a role for mitochondria in regulating synaptic activity (Lee et al. 2018; Picard et al. 2013), raising the possibility that AT-1-induced changes in metabolism could impact neuronal communication and ultimately behavior. The first step in testing this idea is to determine the metabolic and functional consequences of increased AT-1 activity and define how metabolism and neuronal communication are interlinked.

## Results

### AT-1 impacts neuronal metabolism and growth pathways

The endoplasmic reticulum-associated acetyl-CoA transporter AT-1 plays a role in protein quality control in the secretory pathway (**Fig.1A**), but its broader impact on cellular function is less well understood. The impact of AT-1 overexpression (AT-1^sTg^) on mitochondrial and neuronal processes was investigated in primary cortical neurons isolated from embryonic day 17 (E17) wild-type (WT) and AT-1^sTg^ mice. Using an unbiased approach, transcriptional profiles were generated from neurons at day 14 in culture. Bulk RNA-sequencing yielded 800 million sequencing reads (∼130 million per sample). After adapter trimming, raw reads were aligned to the mouse genome (GRCm39), and ∼30,000 genes were quantified. Of these, 632 genes were significantly differentially expressed between genotypes (FDR < 5%) (**Fig.1B** and **S1A**). Among the genes most highly expressed in wild-type neurons were *C3*, a key component of the complement system; *Gja1 and Gjb2*, gap junction proteins; and *Gpam*, a mitochondrial glycerol-3-phosphate acyltransferase. Genes more highly expressed in the AT-1^sTg^ neurons included *Galtn3*, which enables polypeptide N-acetylgalactosaminyltransferase activity, *Cers6*, a ceramide synthase, and *Csrnp3*, a cysteine-serine-rich nuclear protein that enables DNA-binding transcription activator activity. Gene set enrichment analysis 1/19/2026 2:56:00 PM identified 662 pathways in the AT-1^sTg^ dataset (FDR < 5%), including immune and secretory pathways that were significantly downregulated in AT-1^sTg^ neurons, and pathways relating to metabolism, translation, and synaptic regulation that were upregulated in AT-1^sTg^ neurons (**Fig.1C, Table S1B**). Previous studies in the AT-1^sTg^ model reported a progeroid phenotype, with mice having reduced lifespan, cardiomegaly, muscle atrophy, systemic inflammation, and metabolic impairment (Peng et al. 2018). To determine if the AT-1^sTg^ neurons present an “aged” transcriptional signature, the GSEA Gene Ontology (biological processes, molecular function, and cellular component) output of AT-1^sTg^ versus WT neurons was compared to a previously published cortical aging signature (30 months old versus 10 months old) (Souder et al. 2025) (**Fig.1D**). GSEA of the mouse aging cortex dataset (FDR < 5%, **Table S1C**) identified 1152 pathways, of which 353 pathways were shared (**Table S1D**). The “aged” phenotype includes changes in pathways associated with the ribosome, metabolism, synaptic translation, synapse pruning, and immune-related pathways. In the comparison between neurons and the aged cortex, ribosomal pathways stand out as being shared, concordant, and positively enriched. Shared negatively enriched pathways included amine transport and hormone activity (**Fig.1D**). Perhaps not surprisingly, given the comparison between tissue and isolated cells, the immune system pathways identified as positively enriched with aging were not also enriched in AT-1^sTg^ neurons (**Fig.1D, Table S1D**). These data show that AT-1^sTg^ impacts diverse aspects of neuronal function and partially recapitulates the aging signature of the brain.

**Figure 1.**
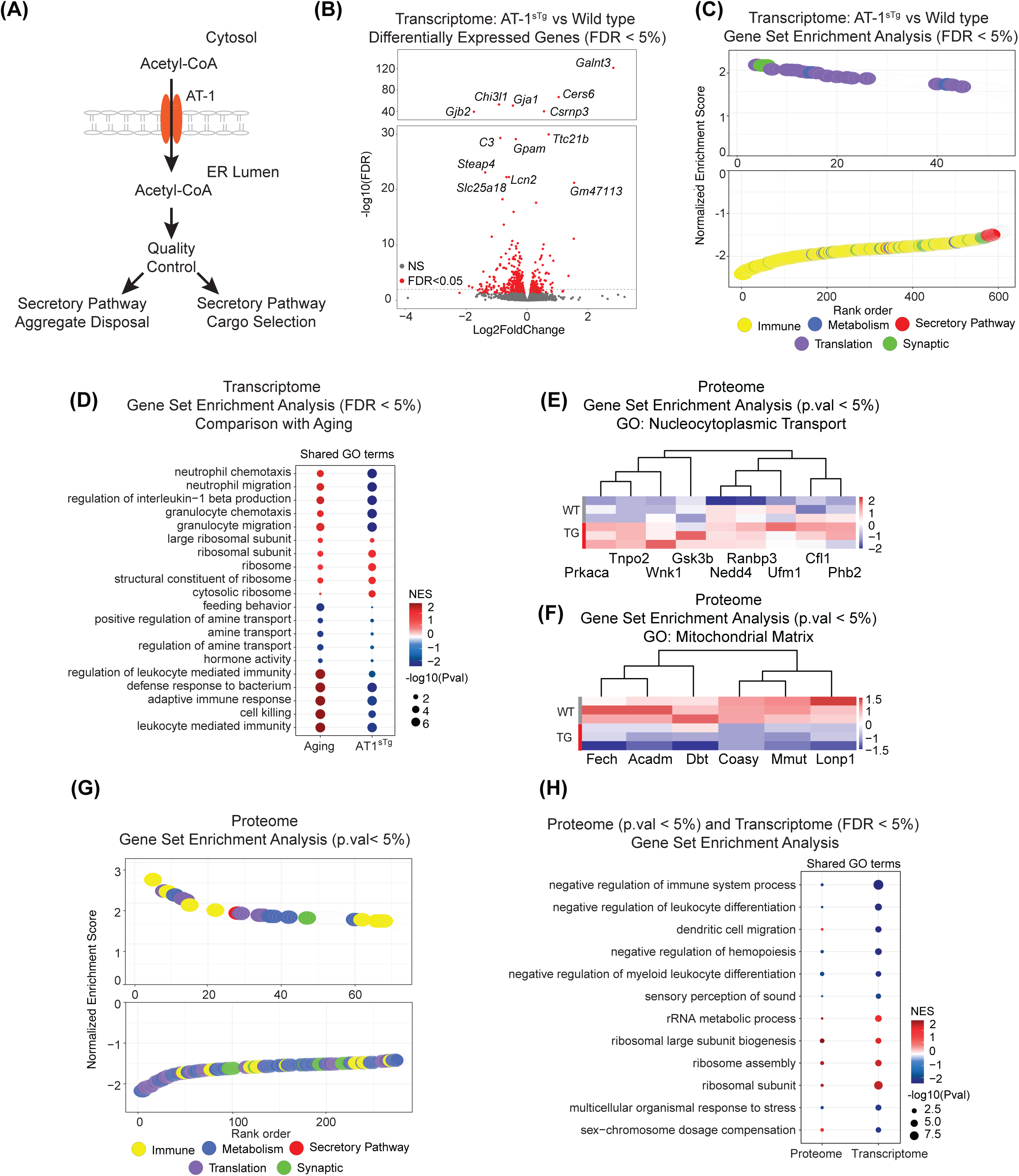
AT-1 impacts neuronal metabolism and growth pathways. A) Schematic of AT-1’s function as an endoplasmic reticulum (ER)-resident acetyl-CoA transporter. Once inside the ER, acetyl-CoA is used as a substrate to acetylate properly folded proteins in the secretory pathway. B) Volcano plot displaying genes quantified via RNA-sequencing. Statistically significant genes are highlighted in red in AT-1^sTg^ primary neurons compared to WT. C) Rank order plot of enriched GO terms in RNA-sequencing dataset by GSEA. Plot is ranked by normalized enrichment score. Points are color-coded by category: immune – yellow, metabolism – blue, secretory pathway – red, translation – purple, synaptic – green. D) Dot plot of shared GO terms enriched in AT-1^sTg^ primary neurons and aging mouse cortex (GSE246478). E) Heatmap of nucleocytoplasmic transport pathway detected by GSEA in AT-1^sTg^ primary neuron proteomics compared to WT. F) Heatmap of mitochondrial matrix pathway detected by GSEA in AT-1^sTg^ primary neuron proteomics compared to WT. G) Rank order plot of enriched GO terms in proteomics dataset by GSEA. Plot is ranked by normalized enrichment score. Points are color-coded by category: immune – yellow, metabolism – blue, secretory pathway – red, translation – purple, synaptic – green. H) Dot plot of shared GO terms enriched in AT-1^sTg^ primary neuron RNA-sequencing and proteomics.

Growing evidence indicates that not only do transcriptional changes fail to predict proteomic changes, but often concordance between the two methods is low (Wang et al. 2019; Maier et al. 2009). The possibility that the lack of accord could be a sampling problem is overcome by using the exact same isolates for RNA-seq and proteomic analysis. Proteins were isolated and proteomic profiles generated via liquid chromatography-tandem mass spectrometry (LC-MS/MS). The detected proteome included 2434 quantified proteins, of which 2413 matched to unique genes (**Table S1E**). The proteomics dataset revealed only two proteins with differential abundance passing significance at FDR < 5%: TNPO2, involved in nuclear import (**Fig.1E, Table S1E**), and COASY, the enzyme CoA synthase, and without multiple testing adjustment, this number increased to 74 proteins (**Fig.1F, Table S1E**). GSEA identified 349 enriched pathways at a nominal p-value < 5% and revealed a similar pattern to that seen in the RNA-seq GSEA. Specifically, the genotype-dependent proteome differences included pathways involved in metabolism, translation, synaptic function, and secretion (**Fig.1G**, **Table S1F**). Interestingly, fewer immune-related pathways were found in the proteome analysis than in the RNA-seq, although the direction of impact was congruent. The proteomics GSEA yielded 407 pathways with p-values < 5%, of which 15 were common between the RNA-seq and proteome (**Fig. 1H, Table S1G**). The positive enrichment of ribosomal pathways at protein and transcript levels stands out among the shared signatures. Other shared pathways of interest included negative enrichment of development and differentiation pathways. These data show that in AT-1^sTg^ neurons, metabolism and translation pathways are engaged at the transcriptional and proteomic levels, and that metabolism and growth-associated pathways communicate, in part, through acetyl-CoA availability.

### AT-1 overexpression drives increased neuronal mitochondrial activity

Cytosolic acetyl-CoA pools are maintained through multiple processes, including lipolysis and the conversion of citrate to acetyl-CoA by ACLY (Shi & Tu 2015). In mitotic cells and neurons, increased AT-1 expression promotes acetyl-CoA transport from the cytosol to the ER lumen (Jonas et al. 2010), but the consequences of this shift on mitochondria have not been characterized. Peroxisome proliferator-activated receptor-gamma coactivator 1-alpha (PGC-1α) is a transcriptional coactivator and master regulator of mitochondrial function (Miller, Clark & Anderson 2019). The brain contains multiple PGC-1α isoforms, PGC-1α1, PGC-1α4, and PGC-1α B1E2, that are expressed from three promoter regions (Soyal et al. 2020). In primary neurons, PGC-1α B1E2 is the most abundant and functionally dominant isoform as far as mitochondrial activity regulation is concerned (Souder et al. 2025). As expected, PGC-1α B1E2 was the most abundant isoform for both WT and AT-1^sTg^ neurons (**Fig.2A**). When compared to WT expression levels, AT-1^sTg^ neurons showed significant increases in both PGC-1α1 and PGC-1α B1E2, and a non-significant trend for increased PGC-1α4 (**Fig.2B**). Expression of known PGC-1α gene targets was increased in AT-1^sTg^ neurons with an overall effect of genotype, and expression of oxidative phosphorylation genes *Cox5b* and *Cycs,* TCA cycle gene *Idh3a,* and antioxidant gene *Sod2* were significantly increased (**Fig.2C**). Unexpectedly, pyruvate dehydrogenase kinase (*Pdk4*), which inhibits glucose influx into the TCA cycle by inhibiting pyruvate conversion to acetyl-CoA, was also upregulated (**Fig.2C**). An impact of AT-1^sTg^ on mitochondrial processes was further corroborated in proteomic profiles, where pathways of mitochondrial fusion, mitochondrial protein targeting, mitochondrial outer membrane and matrix processes were all enriched in AT-1^sTg^ primary cortical neurons (**Fig.2D**).

**Figure 2.**
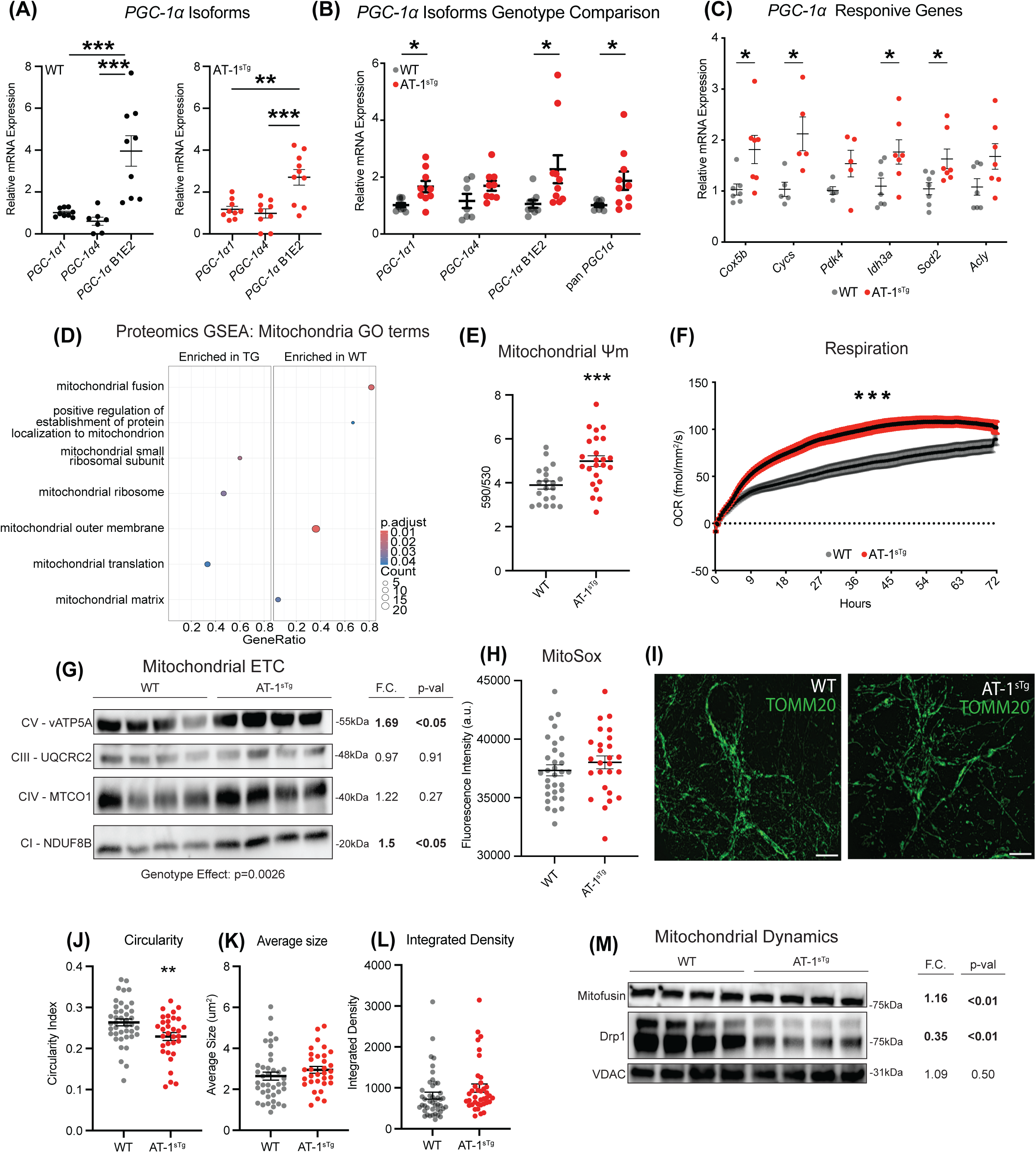
AT-1 overexpression increases neuronal mitochondrial activity. A) RT-qPCR detection of Ppargc1a (PGC-1α) transcript variants in WT and AT-1^sTg^ primary neurons (n=7-10). Data are normalized to the expression of PGC-1α1 in each genotype. B) RT-qPCR detection of PGC-1α transcript variants in AT-1^sTg^ primary neurons (n=7-10). Expression level of each isoform is normalized to WT expression. C) RT-qPCR detection of PGC-1α-responsive genes detected in primary neurons (n=5-7). D) Mitochondrial GO terms identified in GSEA of proteomics from WT and AT-1^sTg^ primary neurons. E) Mitochondrial membrane potential assessed by JC-1 assay in primary neurons (n=20-24). F) Oxygen consumption of primary neurons measured by RESIPHER monitor for 72 hours (DIV4-7). G) Immunodetection of electron transport chain proteins in WT and AT-1^sTg^ primary neurons (n=4 neuron lysates). H) Mitochondrial reactive oxygen species detected by MitoSOX red assay (n=26-32). I) Representative images of mitochondria detected with TOMM20 antibody. J-L) Mitochondrial morphology was determined through analysis in ImageJ to detect mitochondrial circularity (J) size (K) and integrated density (L) (n=38-43 neurons). M) Immunodetection of mitochondrial dynamic regulators Mitofusin, DRP1, and voltage dependent anion channel (VDAC) by Western blot. Data shown as mean ± SEM. (*:p<0.05, **:p<0.01, ***:p<0.001).

Mitochondrial membrane potential measured via JC-1 staining was higher in AT-1^sTg^ neurons (**Fig.2E**), and cellular oxygen consumption rate measured in situ in tissue culture was also elevated compared to WT (**Fig.2F**). The expression of mitochondrial electron transport chain proteins was also altered in AT-1^sTg^ neurons, with a main effect of genotype. Complex I and complex V were significantly higher in the AT-1^sTg^ neurons, with complex IV trending higher and complex III unchanged (**Fig.2G**). These changes in the electron transport chain protein content aligned with the functional outcome of increased cellular respiration. Reactive oxygen species (ROS) detected via MitoSox were not different between AT-1^sTg^ neurons and WT (**Fig.2H**), suggesting that even though respiration was higher, there was no burden of ROS. It is unclear if this is due to unchanged ROS production or to greater clearance of ROS as might be suggested by the induction of *Sod2*.

Mitochondrial morphology, assessed via immunofluorescence (IF) of the mitochondrial membrane protein TOMM20, was also altered in AT-1^sTg^ neurons (**Fig. 2I**). Quantitative analysis revealed a more reticulated mitochondrial network, with significantly lower mitochondrial circularity (**Fig.2J**). In general, this arrangement of mitochondrial networks is associated with greater respiratory capacity (Picard et al. 2013). Changes in mitochondrial function induced in the AT-1^sTg^ neurons were not associated with overt differences in mitochondrial size (**Fig.2K**) or integrated density (**Fig.2L**). Although these were steady-state measures, there was also a suggestion that mitochondrial dynamics may be responsive to altered acetyl-CoA distribution. Western blot detection revealed that fusion-related Mitofusin2 (MFN2) was significantly elevated in AT-1^sTg^ neurons compared to WT, while fission-related Dynamin-Related Protein 1 (DRP1) was significantly lower in AT-1^sTg^ neurons (**Fig.2M**). Consistent with the IF mitochondrial density data, the western blot for voltage-dependent anion channel (VDAC) expression, a surrogate for mitochondrial density, was also equivalent between genotypes (**Fig. 2M**). These data show that intracellular acetyl-CoA availability directs metabolic adaptation, inducing a more fused mitochondrial network with higher membrane potential and respiratory activity.

### AT-1 overexpression alters the fate of cellular glucose

Several of the genes identified as upregulated in response to AT-1 overexpression in primary neurons are associated with the Tricarboxylic acid (TCA) cycle and electron transport chain (ETC), suggesting increased respiration, but the significant change in *Pdk4* would appear to oppose glucose-derived pyruvate entering the TCA. These transcriptional changes suggest a potential difference in fuel substrate utilization, particularly a lowering of glucose utilization for respiration. Intracellular glucose has three potential fates once it becomes phosphorylated: glycolysis, pentose phosphate pathway (PPP, which produces precursors for nucleotide turnover), or storage as glycogen (**Fig.3A**). Differences in glucose handling were suggested in the proteome data, with one of the significant GO terms from the GSEA, Glucose-6-phosphate metabolic process, displaying decreased expression in AT-1^sTg^ neurons (**Fig.3B**). Metabolite tracing in WT and AT-1^sTg^ neurons using uniformly 13C-labeled glucose (U13C; all 6 carbons labeled), and labeling patterns of glucose derived metabolites were quantified by LC-MS/MS. There were no detectable differences in U13C labeling of TCA cycle metabolites between the genotypes (**Fig.3C**). Intracellular lactate detected via enzyme-linked assay was not different between the genotypes (**Fig.3D**), nor were differences detected in extracellular lactate released from neurons.

**Figure 3.**
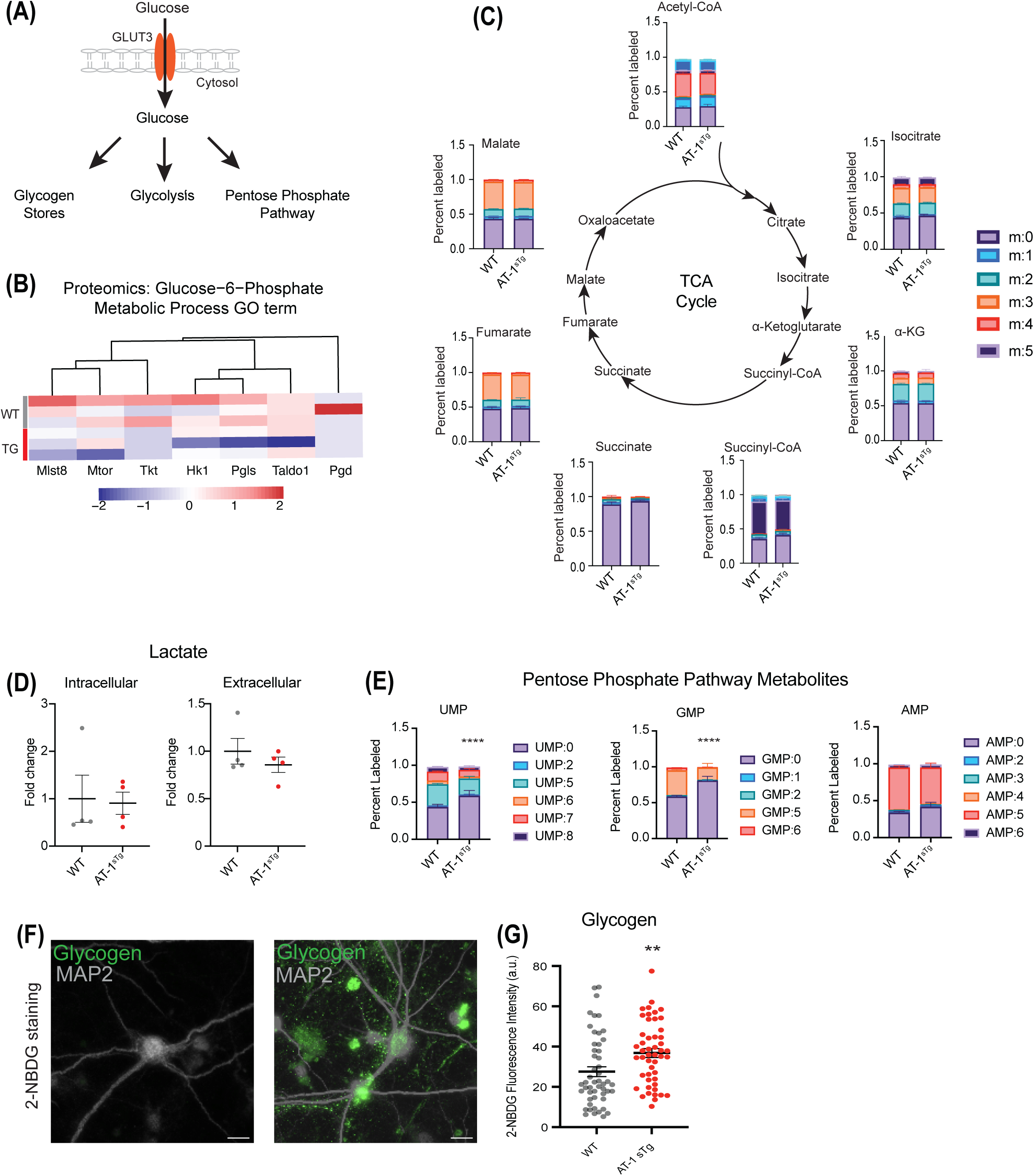
AT-1^sTg^ neurons shuttle glucose toward glycogen storage and away from the pentose phosphate pathway. A) Schematic showing the three fates of glucose once glucose enters a neuron through GLUT3. B) Heatmap of proteins detected in the Glucose-6-phosphate metabolic GO term. C) Schematic showing the incorporation of U^13C^-glucose into the TCA cycle. D) Intracellular and extracellular lactate levels detected in WT and AT-1^sTg^ neurons (n=4). E) Labeling pattern of pentose phosphate metabolites UMP, GMP, and AMP. Bar graph represents the mean across 3-5 independent samples. F-G) Representative images (F) and quantification (G) of glycogen stores detected by the incorporation of fluorescent glucose analog 2-NBDG after 6 hours of labeling (n=49-51 neurons). Data shown as mean ± SEM. (**:p<0.01, ****:p<0.0001).

There was a significant decrease in the labeling of PPP products GMP and UMP (**Fig.3E**). The lack of difference in TCA and reduced labeling of PPP metabolites suggested that glucose may be diverted to stores in AT-1^sTg^ neurons. In general, neurons do not store glycogen at high levels under basal conditions, but aberrant neuronal glycogen has been reported in neurodegenerative disease (Bar et al. 2025; Rai et al. 2018) and hypoxia (Saez et al. 2014). Glycogen storage in WT and AT-1^sTg^ neurons was assessed via 2-NBDG, a fluorescent glucose analog that detects glycogen stores (Zhu et al. 2020). Glycogen storage was strikingly enhanced in AT-1^sTg^ neurons (**Fig.3F**), and quantitative analysis confirmed significant differences between genotypes (**Fig.3G**). These data suggest that the fate of glucose can be somehow influenced by acetyl-CoA-directed signaling. One potential explanation is that the increased respiration observed in the AT-1^sTg^ neurons is due to greater use of non-glucose substrates that could conceivably offset the increased traffic of acetyl-CoA into the ER.

### AT-1 overexpression alters fuel substrate flexibility and lipid storage

Increased expression of PGC-1α and its target genes involved in lipid metabolism, along with increased abundance of ETC components in neurons, suggests that lipid fuel could be a source for energy derivation in AT-1^sTg^ neurons. If so, a genotype effect on lipid stores might be expected. Lipid droplets were detected in primary neurons using LipidTox, a fluorescent dye that binds neutral lipid stores (**Fig.4A**). AT-1^sTg^ neurons had significantly fewer and smaller lipid droplets than WT neurons and total lipid area was lower compared to that of WT neurons (**Fig.4B**). These data indicate that AT-1^sTg^ neurons are not storing lipids, but it is unclear if this is a deficit in synthesis and storage or an increase in depletion due to reliance on lipids as fuel. Fully mature AT-1^sTg^ and WT neurons were treated with a series of inhibitors and monitored for cellular respiration to assess fuel substrate use and flexibility (**Fig.4C**). Three challenges were undertaken: UK5099 (10*μ*M) to block glucose utilization, BPTES (20*μ*M) to inhibit glutaminase, or Etomoxir to inhibit the transport of cytosolic lipids into the mitochondria for oxidation. Respiration dropped acutely in UK5099 treated neurons, with recovery in ∼2 hours and no impact of genotype (**Fig.4D**). Respiration was lower in BPTES treated neurons but with a greater impact in WT than AT-1^sTg^ and with return to baseline oxygen consumption faster in AT-1^sTg^ than WT neurons (**Fig.4E**). The impact of etomoxir on respiration was greater in WT than AT-1^sTg^ neurons and again return to baseline oxygen consumption faster in AT-1^sTg^ (**Fig.4F**). These data confirm that pyruvate is a primary source of fuel in neurons and indicate that AT-1^sTg^ neurons are better able to tolerate deficiency in glutaminase and lipid energy derivation pathways. Perhaps the increased availability of glycogen stores in the AT-1^sTg^ neurons enables a more rapid fuel switch when mitochondrial lipid and amino acid transport are blocked. The fact that AT-1^sTg^ neurons are refractory to etomoxir suggests a storage problem rather than an increased depletion of lipid droplet stores.

**Figure 4.**
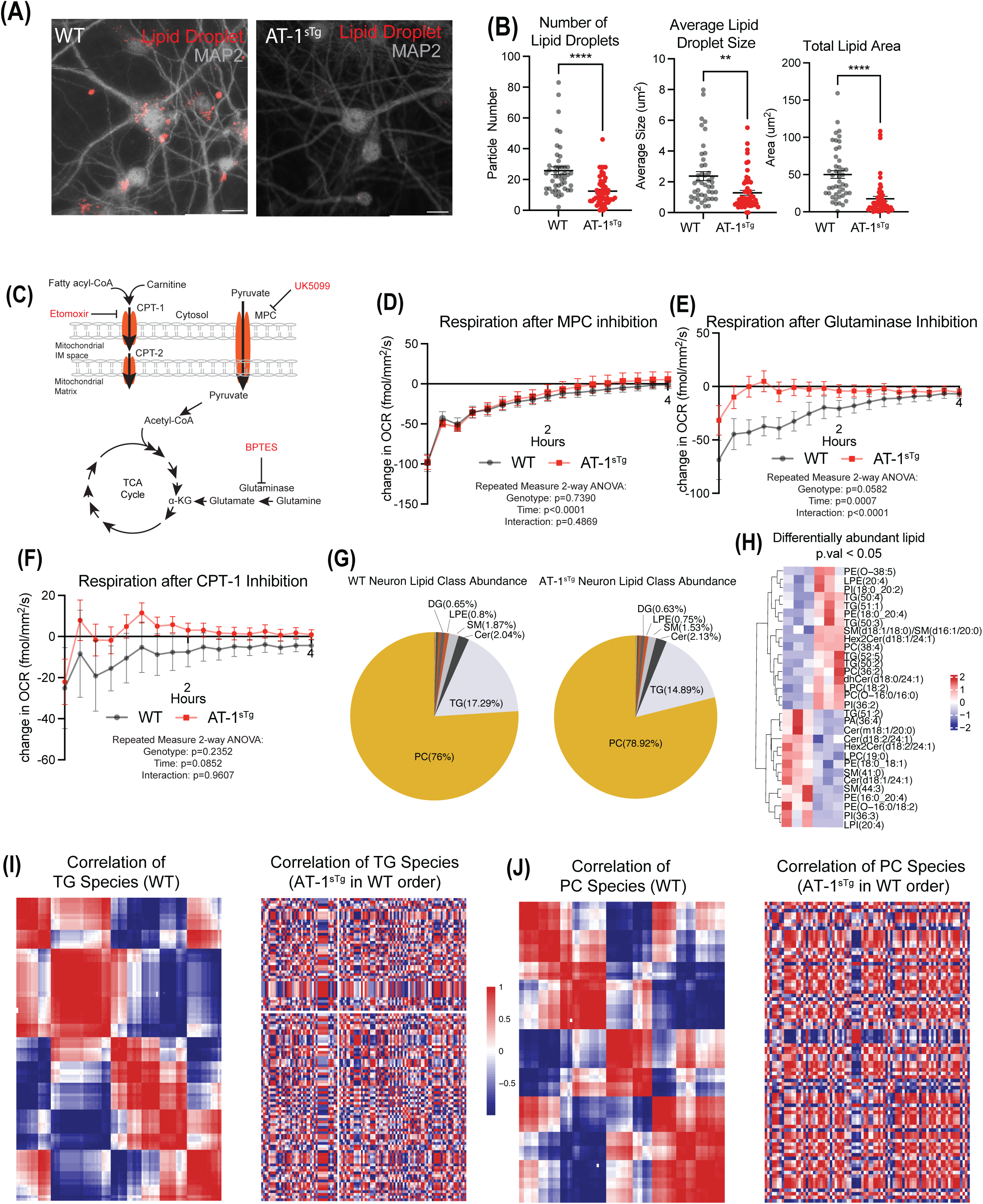
AT-1^sTg^ neurons have altered fuel substrate flexibility. A) Representative images of lipid droplet detection determined by LipidTox incorporation. B) Quantification of lipid droplet number, size, and total lipid area (n=45-56 neurons). C) Schematic of mitochondrial fuel substrate utilization with inhibitors of fatty acid utilization (Etomoxir), pyruvate influx (UK5099), or amino acid utilization (BPTES). D-F) Change in oxygen consumption of primary neurons treated with inhibitors of mitochondrial glucose (D) (n=5-10), amino acid (E) (n=9-17), or lipid (F) (n=9-17), or utilization. G) Pie charts showing the distribution of lipid species detected via LC-MS in WT and AT-1^sTg^ neurons. H) Heatmap of 31 differentially abundant lipids (p-value < 0.05). I) Pearson rank correlations of triglycerides in WT (left) and AT-1^sTg^ neurons (right) following hierarchical clustering. Both maps are ordered using the clustering of WT triglycerides. J) Pearson rank correlations of phosphatidylcholines in WT (left) and A AT-1^sTg^ neurons (right) following hierarchical clustering. Both maps are ordered using the clustering of WT phosphatidylcholines. Data shown as mean ± SEM. (**:p<0.01, ****:p<0.0001).

The overt lipid droplet phenotype suggested that lipid metabolism might be altered in the AT-1^sTg^ neurons. The impact of AT-1^sTg^ on the neuron lipidome was investigated via global targeted mass spectrometric lipidomics. This method is set up to detect 750 lipids and lipid extracts from these neurons yielded 448 lipid species. The distribution of lipid classes was not significantly different by genotype, with the most abundant classes being phosphatidylcholines (PC) and triacylglycerides (TG) in both genotypes (**Fig.4G**). Of the 448 lipids detected using this targeted approach, only 31 lipids were differentially abundant between WT and AT-1^sTg^ neurons (p.val < 0.05, **Fig.4H**, **Table S1H**), including a mix of species of triglycerides, ceramides, and ceramide derivatives, among others.

Lipids are a highly complex class of molecules with interconversions such as desaturation and elongation occurring across classes, in addition to connections among synthetic pathways, resulting in linked abundance and composition by class. To interrogate interactions among species within and among lipid classes, Pearson rank correlations were calculated and visualized using color-coded correlation values (red: positive; blue: negative). Even though lipid composition was not different by genotype, associations among species both within and among lipid classes were entirely disrupted in the AT-1^sTg^ neurons (**Fig.S1**). When the AT-1^sTg^ neuron lipidome is clustered by the order determined by the wild-type lipidome, there are no consistencies between the genotypes. This is also seen in the reverse comparison: when the wild-type lipidome is clustered according to the order determined by the AT-1^sTg^ lipidome, there is no concordance. Focusing on the most abundant species, hierarchical clustering of correlations among lipid species of TG revealed a series of highly correlated modules in WT neurons (**Fig.4I**). When constraining lipid order using WT clustering and applying the AT-1^sTg^ lipidome, cohesive modules of interrelated species were no longer detected. Patterns of correlation were completely altered, indicating that processing within the TG lipid class is distinct for AT-1^sTg^ neurons. The same analysis approach for PC revealed a similar outcome, where discrete clusters of highly correlated species were detected in WT neurons but were entirely absent in AT-1^sTg^ neurons (**Fig. 4J**). Even among less abundant species such as diacylglycerides and ceramides, clear patterns of correlation in WT are altered in the AT-1^sTg^ neuronal lipidome. Although these data cannot explain differences in lipid storage among genotypes, acetyl-CoA availability appears to have a major influence on fundamental lipid processing in neurons that is manifest across diverse lipid classes.

### AT-1 overexpression alters redox metabolism and the metabolic microenvironment

Even modest changes in mitochondrial oxidative capacity precipitate changes in nuclear and cytosolic redox metabolism (Miller, Clark, Martin, et al. 2019), indicating that cellular redox is matched to prevailing mitochondrial energetic status. Mitochondrial activity, as well as many other catabolic and anabolic processes, are dependent on the redox cofactors NAD and NADP. Proteomics identified NADH metabolic processes as negatively enriched in the AT-1^sTg^ neurons, raising the possibility that redox might also be altered in AT-1^sTg^ neurons (**Fig.5A**). The reduced forms, NADH and NADPH, are autofluorescent (excitation 740nm, emission 450nm), a property that is harnessed in multiphoton laser scanning microscopy (MPLSM) (Schaefer et al. 2019). Fluorescence intensity measures the overall abundance of endogenous NAD(P)H. In neurons, NAD(P)H intensity is dependent on subcellular location, with lower values detected in the nucleus and extremities compared to the extranuclear region of the soma (**Fig.5B**). Cytosolic and nuclear NAD(P)H intensity was higher in AT-1^sTg^ neurons compared to WT, confirming that redox metabolism is sensitive to changes in AT-1. Fluorescence lifetime imaging microscopy (FLIM) quantifies the kinetics of photon release and is distinct for protein-bound and unbound fluorophores, although other factors such as microenvironment and pH may also play a role (Schaefer et al. 2019). The mean fluorescence lifetime (*τ*_m_) can be described by a first-order decay curve ( *τ*_m_ = *τ*_l_a_l_ + *τ*_2_a_2_) (**Fig.5C**) where *τ*_1_ is the short lifetime component which represents the free NAD(P)H in a cell, and *τ*_2_ is the long lifetime component which is protein-bound NAD(P)H. Nuclear *τ*_m_ was lower than cytosolic *τ*_m_ and main effects of subcellular region and of genotype were detected, with higher values in AT-1^sTg^ neurons (**Fig.5D**, **5E**). There was a trend for the relative contribution of free NAD(P)H (a_1_) to be lower in the AT-1^sTg^ neurons, a quality that usually tracks with reliance on glycolytic energy derivation (**Fig.5F**). In AT-1^sTg^ neurons values of both *τ*_1_ (free) and *τ*_2_ (bound) were significantly higher. Several non-metabolic processes are also linked to redox status, including factors that influence gene expression by modulating transcription factors or chromatin. The NAD^+^-dependent protein deacetylase SIRT1 has been associated with the regulation of metabolism, both by regulation of enzymatic function and by regulation of transcription factors that regulate metabolic genes (Chang & Guarente 2014). A significant increase in SIRT1 protein abundance via western blot, along with changes in redox indices, suggested that SIRT1 activity might be altered in AT-1^sTg^ neurons (**Fig.5G**). Consistent with this, a decrease in total cellular acetylated lysine was detected in the AT-1^sTg^ neurons (**Fig.5H**). The transcriptional corepressor CtBP2 is responsive to NAD/NADH (Sekiya et al. 2021). Although a link between CtBP2 expression and redox has not been established, at the protein level, in AT-1^sTg^ neurons, CtBP2 was increased (**Fig.5I**). At the protein level, pathways associated with chromatin remodeling were enriched in the AT-1^sTg^ neurons (**Fig.5J**). Overall, the AT-1^sTg^ neurons are metabolically distinct in terms of redox metabolism, with consequences for post-translational modification via acetylation and potential chromatin remodeling via CtBP2 and others.

**Figure 5.**
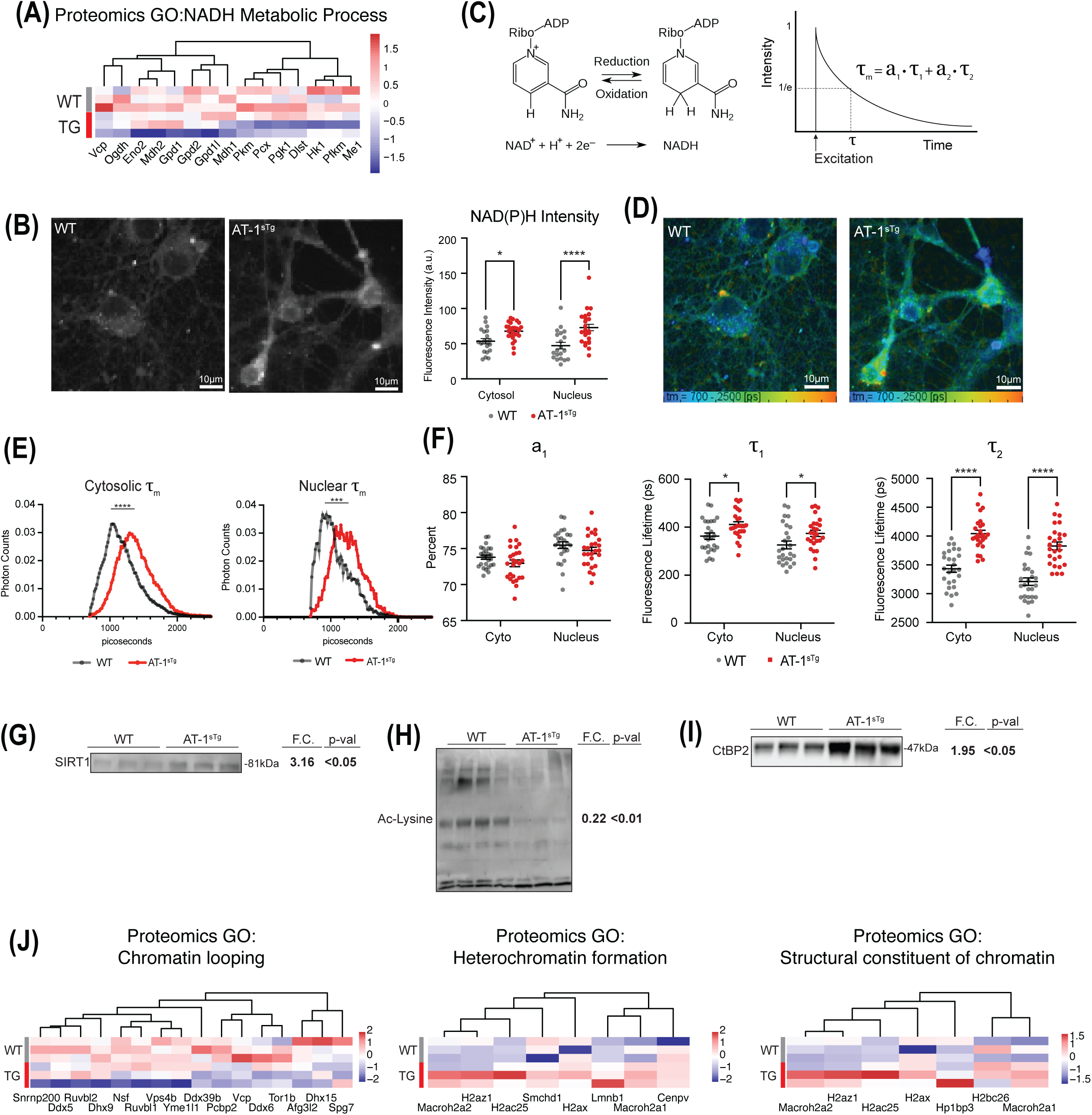
AT-1 overexpression shifts neuronal redox metabolism. A) Heatmap of NADH Metabolic Process detected as significantly changed in proteomics GSEA. B) Representative images and quantification of NAD(P)H fluorescence intensity detected by multiphoton laser scanning microscopy. C) Schematic of NAD and NADH reduction and fluorescence lifetime decay curve. D-E) Representative images (D) and distributions (E) of NAD(P)H mean fluorescence lifetime of WT and AT-1^sTg^ neurons (n=26-27 neurons). F) Quantitation of the relative contribution of free NAD(P)H (a_1_) to the mean fluorescence lifetime, the short component of the decay curve (*τ*_1_), and the long component of the decay curve(*τ*_2_). G) Immunodetection of SIRT1 in primary neurons (n=4). H) Immunodetection of acetylated lysine in primary WT and AT-1^sTg^ neurons (n=4 WT, n=3 AT-1^sTg^). I) Immunodetection of CtBP2 primary WT and AT-1^sTg^ neurons (n=4 WT, n=3 AT-1^sTg^). J) Heatmap of chromatin looping, heterochromatin formation, and structural constituent of chromatin detected as significantly changed in proteomics GSEA. Data shown as mean ± SEM. (*:p<0.05, ***:p<0.001, ****:p<0.0001).

### AT-1 overexpression disrupts synaptic communication

In primary neurons, differences in mitochondrial function are associated with changes in soma size, arborization, and electrophysiological properties (McGregor et al. 2025; Souder et al. 2025). Neuron-specific overexpression of AT-1 increases neuronal arborization in the mouse hippocampus, and a connection with mitochondrial function was hinted at in the hippocampal proteome (Hullinger et al. 2016). The possibility that changes in AT-1^sTg^ neuronal metabolism described above could have consequences for cell morphology and dendritic architecture was explored using Scholl analysis. AT-1^sTg^ neurons had increased numbers of branches compared to WT (**Fig.6A**). Although specific to neuron type, in cortical neurons, the degree of arborization can sometimes be linked to soma size (Benavides-Piccione et al. 2024). This was not evident in AT-1^sTg^ neurons. This suggests that structural features known to be linked to metabolism are contributing to the expanded area of dendritic contact. Consistent with this idea, RNA-seq GSEA analysis revealed significant enrichment of cell adhesion, filament, and integrin pathway GO terms in the AT-1^sTg^ neurons (**Fig.6B**). These cellular structural pathways have been linked to neurite outgrowth, the formation and maintenance of synapses, and synaptic plasticity (Lilja & Ivaska 2018).

**Figure 6.**
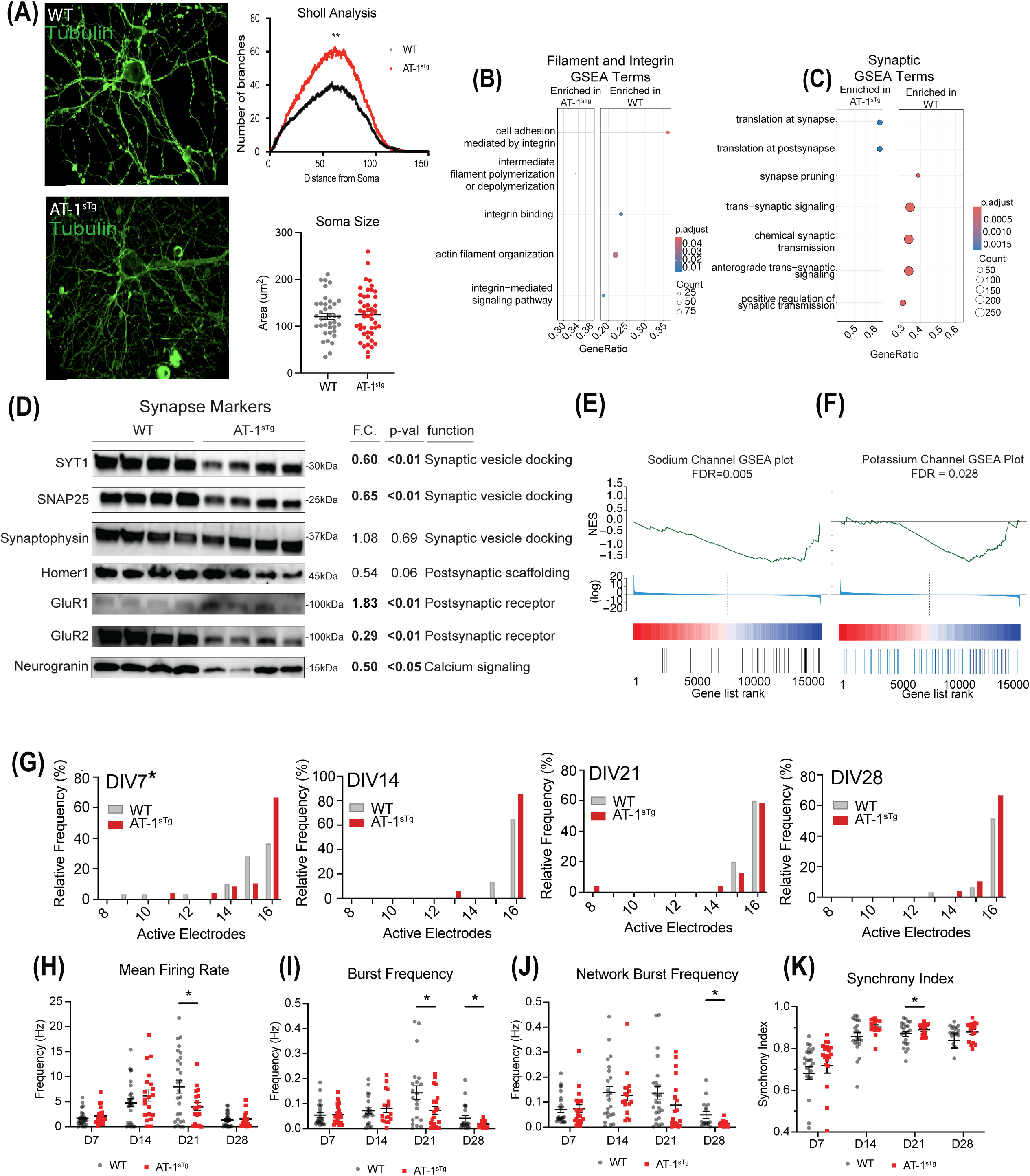
AT-1^sTg^ neurons have altered synaptic communication. A) Representative images (left) and quantification (right) of neuronal architecture detected by immunodetection of tubulin. Dendritic arborization of WT and AT-1^sTg^ neurons quantified by Sholl analysis. Soma size was quantified in ImageJ (n=38-46 neurons). B-C) Dotplots showing GO terms related to filament and integrin (B) and synaptic processes (C) enriched in the RNA-seq dataset. D) Immunodetection of synaptic proteins in DIV15 WT and AT-1^sTg^ neurons (n=4). E) GSEA plot of sodium channels detected in the RNA-seq data of AT-1^sTg^ neurons compared to WT. E) GSEA plot of potassium channels detected in the RNA-seq data of AT-1^sTg^ neurons compared to WT. G-K) MEA spontaneous activity measured at DIV7, 14, 21, and 28. G) Histograms showing the number of active electrodes per network expressed as the relative frequency in percent. Eight active electrodes were used as the minimum value required for a mature network. G) Spontaneous activity is measured by mean firing rate (H), burst frequency (I), network burst frequency(J), and synchrony index (K). Each data point is an independent network of cultured neurons (n=16-25). MEA data is from five WT and four AT-1^sTg^ mice. Data shown as mean ± SEM. (*:p<0.05, **:p<0.01).

The greater arborization of the AT-1^sTg^ neurons suggested that connectivity among neurons might also be increased. Using an unbiased approach, genotype differences in synaptic-related GO terms were detected in the RNA-seq analysis (**Fig. 6C**). AT-1^sTg^ neurons were enriched for translation at synapses but repressed synaptic pruning and synapse-associated communication processes. At the proteome level, AT-1^sTg^ neurons were negatively enriched in synaptic organization, synapse-associated communication, and axonal transport. These data suggested lower, not greater, synaptic density on the expanded dendrites. Taking a targeted approach, western blot analysis showed that the abundance of markers of synaptic vesicle loading and synaptic vesicle cycle (SYT1, SNAP25) were lower in AT-1^sTg^ neurons (**Fig.6D**). Expression of Synaptophysin and Homer1, proteins that play a role in synaptic formation, was also investigated. While Synaptophysin was unchanged between the genotypes, Homer1 expression was lower in the AT-1^sTg^ neurons. Next, the expression of proteins involved in synaptic communication was investigated. AMPA receptor subunits GluR1 and GluR2 are post-synaptic proteins involved in glutamatergic signaling. Expression of GluR1 was higher in AT-1^sTg^ neurons, while GluR2 was significantly down-regulated. The significance of this shift in GluR subunit balance is unclear; however, changes in GluR subunit balance have been implicated in synaptic plasticity (Zhao et al. 2020).

In mice overexpressing AT-1 only in neurons, long-term potentiation in hippocampal neurons is altered (Hullinger et al. 2016). Expression of ion channels known to contribute to LTP was investigated using a targeted approach with the RNA-seq data. A significant downregulation of both sodium (**Fig.6E**) and potassium channel (**Fig.6F**) GO terms was detected at the transcript level in AT-1^sTg^ neurons, indicating a likely change in the ability to fire action potentials and communicate effectively. Multielectrode array experiments were conducted to assess the electrophysiological function of AT-1^sTg^ neurons. Spontaneous neuronal activity was monitored for 10 minutes in 24-well multielectrode array plates. Activity was measured at four time points during the maturation of neurons (DIV7, 14, 21, and 28) to capture potential key points of developmental differences. The network was deemed mature when more than 50% of the electrodes were active over the course of the recording. AT-1^sTg^ neurons show a higher percentage of mature networks at DIV7 compared to wild-type (**Fig. 6G**), but by DIV14, WT neurons did not differ significantly in the number of active electrodes from AT-1^sTg^ neurons. These data are in line with the outgrowth imaging showing that AT-1^sTg^ neurons have increased dendritic complexity and that this leads to increased synaptic communication in the early stages of maturation. This early apparent advantage was lost by DIV21. In the fully mature network, the number of active electrodes did not differ between genotypes; however, the mean action potential firing rate (**Fig. 6H**), burst frequency (**Fig. 6I**), and network burst frequency (**Fig. 6J**) were significantly lower in AT-1^sTg^ neurons at DIV21. The burst frequency and network burst frequency were lower again in the AT-1^sTg^ neurons at DIV28. A greater degree of firing synchrony in AT-1^sTg^ neurons was suggested by the numerically higher synchrony index across all time points in the experiment (**Fig. 6K**). This indicates that although the AT-1^sTg^ can form a coordinated network, transmission through that network is compromised, with loss of functionality appearing at the later stages of maturation. Overall, these data indicate a tight link between changes in metabolism and neuronal architecture and connect metabolic status to synaptic function.

## Discussion

In this study, we defined the metabolic and synaptic changes that occur in neurons due to AT-1 overexpression, revealing interconnections between the regulation of ER function in the secretory pathway, the well-established role of AT-1, and broader neuronal health and function. In AT-1^sTg^ neurons, mitochondrial membrane potential was increased, and the remodeling of mitochondrial morphology toward filamentous networks was consistent with the increase in cellular respiration. These changes might reasonably be anticipated as a means to restore cytosolic acetyl-CoA, given that AT-1 overexpression diverts acetyl-CoA into the ER. The most obvious sources of acetyl-CoA include derivation from TCA intermediates or as part of fatty acid oxidation. Metabolite tracer studies did not show differences in the TCA cycle under pseudo-steady-state conditions, specifically when glucose is the input. This argues that the increase in respiration is supported by acquired pyruvate (from media or amino acids) or by lipid sources. The AT-1^sTg^ neurons were almost completely deprived of lipid droplet stores, favoring the idea of increased lipid utilization; however, the inhibitor studies suggest that there is a more complex metabolic shift in the AT-1^sTg^ neurons. Specifically, inhibition of CPT-1, the fatty acid transporter for mitochondrial lipid oxidation, had less impact on AT-1^sTg^ neurons than on wildtype, indicating that there is no increase in dependence on lipid fuel.

The accumulation of glycogen in the AT-1^sTg^ neurons was an unexpected phenotype. Although glycogen flux was not included in the tracer studies, the lower flux toward pentose phosphate in the absence of a change in glycolysis suggests that overexpression of AT-1 causes a diversion of glucose toward glycogen storage. In general, expanded glycogen stores are associated with neuronal dysfunction. Patients with glycogen storage disorders, such as Lafora disease, present with epilepsy (Turnbull et al. 2011), and in mouse models of the disorder, impairments in the electrophysiological properties of hippocampal synapses have been reported (Duran et al. 2014). In AT-1^sTg^ neurons, there were distinct differences in synaptic activity throughout the early stages of maturation, with AT-1^sTg^ reaching maturity sooner but presenting relative deficits compared to WT at the later stages. Neuronal morphology was also altered by AT-1 overexpression, with greater dendritic arborization, while soma size was not changed. The loss of functionality of the dendrites might be explained by the lowered expression of many synaptic proteins. One possible explanation for this phenotype is the presence of an activity-independent cue (Kirchner et al. 2023). Together, our data suggest that disturbances in cellular distribution and availability of acetyl-CoA cause dysfunctional synaptic communication by uncoupling synaptic maintenance processes from arborization, leading to an inability to produce proper communication networks.

It is clear that metabolism is altered in the AT-1^sTg^ neurons; the AT-1 over-expressors did not differ from WT in vulnerability to pyruvate transport inhibition, but susceptibility to either glutaminase or CPT-1 inhibition was markedly reduced. The solute carrier for glutamate transport, Slc25a18, was significantly lower in the AT-1 neurons, perhaps lowering dependency on the use of glutamate-to-alpha ketoglutarate funneling into the TCA. The reason that AT-1^sTg^ neurons were refractory to etomoxir is not clear. The lack of lipid droplets and increase in respiration would have suggested increased use, if not dependence, on lipid fuel sources; however, it is possible that lipids are depleted not as fuel but as a source for acetyl-CoA. Consistent with this possibility, mice with AT-1 haploinsufficiency (AT-1^S113R/+^) develop spontaneous steatosis while mice with overexpression of AT-1 (AT-1^sTg^) are resistant to diet-induced steatosis (Dieterich et al. 2019). FLIM suggests that the intercellular redox environment has shifted, reflected in the increase in *τ*_1_ values, and the redox-associated proteome is different, reflected in higher *τ*_2_ values. The lack of significant change in ROS despite the increase in mitochondrial membrane potential, increased abundance of ETC components, and increased cellular respiration may be linked to the increase in mitochondrial fusion and *Sod2* expression. Mitochondrial fusion is known to be associated with a lower level of ROS production, and *Sod2* is a critical antioxidant enzyme (Xu et al. 2025). The increase in NAD(P)H availability in the cytosol and nucleus was associated with differences in NAD-dependent factors. Sirt1 is an NAD-dependent deacetylase with targets in both nuclear and cytosolic compartments. Indeed, whole-cell acetylation levels were lower in the AT-1^sTg^ neurons. CtBP2 is a chromatin-associated repressor, and its activation requires NADH-dependent homodimerization (Thio et al. 2004). In the liver, CtBP2 regulates lipid biosynthesis indirectly via Srebp1 (Sekiya et al. 2021). The 2-fold higher abundance of the protein in AT-1^sTg^ neurons could be a contributing factor in the depletion of lipid droplets, where consumption of lipid to restore acetyl-CoA is not sufficiently countered by de novo lipogenesis.

The data shown here highlight the importance of understanding intracellular metabolite sharing and communication networks. In particular, the impact of changes in acetyl-CoA is evident across energetic and synthetic pathways, from mitochondrial-dependent and independent pathways of metabolism to cellular redox and metabolic flexibility. In the context of neurons specifically, the morphological and synaptic processes that are vital to neuronal communication and functional network assembly are sensitive to metabolism. This study underscores the importance of maintaining the integrity of metabolism and uncovers a central role of acetyl-CoA as a sensor and regulator of neuronal health.

## Materials and methods

### Cell culture

Primary cortical neurons were isolated from embryonic day 17 (e17) mice. The cortices were harvested from pups into ice-cold HBSS (Thermo Fisher; 14-175-095; Waltham, MA) where the midbrain and meninges were removed. Brain tissue was minced and digested in 0.25% trypsin (25-200-056; Fisher Scientific; Waltham, MA) for 20 minutes at 37*°*C. Trypsin was quenched with DMEM/10%FBS/1%Pen/Strep, and cells were dissociated and counted prior to plating on poly-d-lysine coated plates. Cultures were then fully changed to Neurobasal Plus Media (2% B27^+^, 1% GlutaMax, 1% Pen/Strep) (Thermo Fisher; A3653401; Waltham, MA) 1-2 hours after plating. Media was changed by ½ volume every 3 days until experiments. All primary neuron experiments were conducted between day 10-15 *in vitro* unless otherwise noted.

### qRT-PCR

Cells were lysed with Trizol (Fisher Scientific; Waltham, MA; 15596018) and RNA was isolated using Zymo Research Direct-zol RNA MiniPrep Kit (R2052; Zymo Research; Irvine, CA). RT-qPCR was conducted using iTaq Universal SYBR Green Supermix (1725121, Bio-Rad, Hercules, CA). Primer sequences for all transcripts can be found in **Table S2**.

### Bioassays and Immunofluorescence

#### Multiphoton laser scanning microscopy

Microscopy was conducted as previously described (Souder et al. 2025). Briefly, neurons were grown on poly-d-lysine coated glass coverslips, fixed in 10% formalin (76263-678; VWR; Allentown, PA) and mounted using Fluoromount (F4680; Sigma-Aldrich; St. Louis, MO, USA). Images were captured using a Nikon 60X 1.3apo objective and a 457/40 filter. Images were acquired over a 60-second period. Image analysis was conducted using SPCImage 8.0. (https://www.becker-hickl.com/products/spcimage/).

#### JC-1

Mitochondrial membrane potential was quantified with JC-1 dye (T3168; Invitrogen, Waltham, MA). Neurons were cultured for 10 days and then incubated with 1ug/mL JC-1 dye for 30 minutes. Cells were washed with PBS prior to fluorescence detection. Fluorescence was measured using excitation/emission wavelengths of 535/590 nm and 485/530 nm.

#### Oxygen consumption (OC)

OC was measured using a Resipher oxygen consumption monitor (Lucid Scientific; Atlanta, GA). OC was measured for 72 hours from DIV 4 to 7. Fuel substrate flexibility was determined on DIV10 neurons using 5uM Etomoxir (CPT-1 inhibitor) (E1905; Sigma-Aldrich; St. Louis, MO, USA), 20*μ*M of BPTES (Glutaminase inhibitor) (SML0601; Sigma-Aldrich; St. Louis, MO), or 10*μ*M UK5099 (5048170001; Sigma-Aldrich; St. Louis, MO). AT-1^sTg^ and WT neurons, were treated with Etomoxir, BPTES, or UK5099 on DIV10, and oxygen consumption was monitored for 24 hours. Change in OCR (OCR) was determined by comparing OCR at 15-minute intervals to the basal OCR before inhibitor treatment.

#### Immunofluorescence

Images were acquired on a Leica DMi8 inverted microscope equipped with a HCX PL Fluotar 100X /1.30 oil objective. Images were analyzed using ImageJ (NIH, Wayne Rasband, http://rsb.info.nih.gov/ij/). Cells were grown on glass coverslips coated with poly-d-lysine. Cells were fixed in 10% formalin (245-684; Fisher Scientific; Waltham, MA), for 10 minutes, permeabilized with 0.3% Triton X-100 (T8787; Sigma-Aldrich; St. Louis, MO) in PBS for 1 hour, and incubated with primary antibody cocktail overnight. Primary antibodies used were: Tomm20 (1:84.2; ab56783; Abcam), MAP2 (1:5000; ab5392; Abcam) and alpha-tubulin (1:200; T6199; Sigma-Aldrich). Cells were washed and then incubated with secondary antibodies for 1 hour. Cells were washed 3X before mounting using Fluoromount (F4680; Sigma-Aldrich; St. Louis, MO, USA).

#### Lipid droplet detection

Lipid droplets were detected using HCS LipidTox Red neutral lipid stain (H34476; ThermoFisher; Waltham, MA) according to manufacturer instructions. Fixed neurons were incubated with MAP2 antibody overnight. Coverslips were washed 3X with PBS to remove antibody. Cells were incubated with LipidTox dye (1:200 dilution) for 30 minutes. Cells were washed 3X with PBS and mounted. Images were acquired on a Leica DMi8 inverted microscope equipped with a HCX PL Fluotar 100X /1.30 oil objective. Lipid droplet particle number, size, and total stained area were analyzed for each cell using ImageJ (NIH, Wayne Rasband, http://rsb.info.nih.gov/ij/).

#### Glycogen labeling

Neurons were incubated for 6 hours with 500 *μ*M 2-NBDG (11046; Cayman Chemical Company; Ann Arbor, MI, USA). Neurons were then washed three times with PBS before fixation and mounting. Images were acquired on a Leica DMi8 inverted microscope equipped with a HCX PL Fluotar 100X /1.30 oil objective. Mean intensity of 2-NBDG was analyzed for each cell using ImageJ (NIH, Wayne Rasband, http://rsb.info.nih.gov/ij/).

#### Sholl analysis

Conducted using the Sholl Analysis plugin for ImageJ.

#### Western Blotting

Cells were lysed, and protein was extracted in a modified RIPA buffer containing protease and phosphatase inhibitors (P8340 and 524624, respectively; Sigma-Aldrich, St. Louis, MO, USA). Proteins were detected by immunoblotting using standard techniques. Western blot densitometry was conducted in ImageJ to detect band intensity. All proteins were normalized to the total protein in the lane detected by Ponceau stain. Antibodies used were:

| Target | Dilution | Manufacturer | Catalog No. |
| --- | --- | --- | --- |
| Total Oxphos Cocktail | 1:250 | Abcam | Ab110413 |
| Mitofusin2 | 1:1000 | Abcam | Ab124773 |
| Drp1 | 1:1000 | Abcam | ab184247 |
| VDAC1 | 1:200 | Abcam | Ab154856 |
| SYT1 | 1:1000 | Abcam | Ab302627 |
| SNAP25 | 1:1000 | Thermo Scientific | MA517609 |
| Synaptophysin | 1:10000 | Abcam | Ab206868 |
| Homer1 | 1:1000 | Synaptic Systems | 160 011 |
| GluR1 | 1:1000 | Thermo Scientific | MA5-27694 |
| GluR2 | 1:250 | Thermo Scientific | 32-0300 |
| Neurogranin | 1:1000 | Abcam | Ab217672 |
| SIRT1 | 1:1000 | Abcam | ab110304 |
| Ac-Lysine | 1:1000 | Cell Signaling Technologies | 9441S |
| CtBP2 | 1:500 | Cell Signaling Technologies | 13256S |

#### Intracellular metabolite extraction and liquid chromatography-mass spectrometry

For glucose tracing experiments, media without D-glucose was supplemented with U-[13C]-D-glucose (CLM-1396-1; Cambridge Isotope Laboratories, Tewksbury, MA) at formulation concentration and was used in the place of chemically identical regular unlabeled Neurobasal media. On day 10 *in vitro*, primary neurons were cultured with a stable isotope for 24 hours with additional media change at 2-h prior to metabolite extraction. Cells were washed three times with Dulbecco’s phosphate-buffered saline (D-PBS) (20-031; Corning; Corning, NY) and metabolites were extracted with LC-MS grade 80:20 methanol and water (v/v). Samples were dried under nitrogen flow and resuspended in LC-MS grade water.

Samples were analyzed using a Thermo Q-Exactive mass spectrometer coupled to a Vanquish Horizon UHPLC. Analytes were separated on a 100 × 2.1 mm, 1.7 µM Acquity UPLC BEH C18 Column (Waters), with a 0.2 ml min−1 flow rate and with a gradient of solvent A (97:3 H2O/methanol, 10 mM TBA, 9 mM acetate, pH 8.2) and solvent B (100% methanol). The gradient was as follows: 0 min, 5% B; 2.5 min, 5% B; 17 min, 95% B; 21 min, 95% B; 21.5 min, 5% B. Data were collected in full-scan negative mode. The setting for the ion source was as follows: 10 aux gas flow rate, 35 sheath gas flow rate, 2 sweep gas flow rate, 3.2 kV spray voltage, 320°C capillary temperature and 300°C heater temperature.

The intracellular metabolites reported were identified based on exact m/z and retention times determined with chemical standards. Data were analyzed with MAVEN (Clasquin et al. 2012; Melamud et al. 2010). Metabolite abundance measured by ion count in LC-MS analysis was normalized to total protein content.

#### RNA isolation and RNA-seq library preparation

Total RNA was extracted using the Direct-zol RNA Kit (Zymo Research) following the manufacturer’s instructions. RNA integrity was assessed with an Agilent 2100 Bioanalyzer. mRNA was isolated from purified total RNA using the NEBNext Poly(A) Magnetic Isolation Module (NEB #E7490), and libraries were prepared using the NEBNext Ultra II RNA Library Prep Kit for Illumina (NEB #E7770). Library quality and concentration were evaluated using the Agilent DNA 1000 Kit and the Qubit High Sensitivity Assay Kit (Invitrogen), respectively. Sequencing was performed on an Illumina NovaSeq platform with 150 bp paired-end reads (∼115 million reads per library).

#### RNA-seq bioinformatics analysis

Libraries were sequenced 2x150bp on an Illumina NovaSeq X Plus with an average of 114M reads per sample. Alighments to the mm39 reference genome and feature counting was performed using Rsubread (v2.18.0) (Liao et al. 2019).

Genes were excluded from downstream analysis if they did not have mapped reads in at least half of the RNA-seq libraries. Surrogate variable analysis (sva) was used to estimate and correct for experimental noise thorugh R package ‘sva’ v. 3.56.0. The ‘removeBatchEffect’ function from ‘limma’ v. 3.64.3 was used to regress out effects of the surrogate variables from raw read counts. The ‘DESeq2’ R package (DESeq2 v.1.48.2) was used to perform differential expression analysis. Significant differentially expressed genes (FDR < 5%) are reported in **Supplementary Data 1A**.

#### Protein extraction and proteomics

Protein pellets from TRIzol extractions were washed with cold acetone and air dried to completion. Dried pellets were re-solubilized in 65µl of 8M Urea in 50mM NH_4_HCO_3_ (pH8.5) and 5µl was taken for BCA protein measurement. Subsequently 30µg of protein in 20µl 8M Urea and 50mM NH_4_HCO_3_ (pH8.5) was taken for tryptic/LysC digestion where the samples were first diluted with 2.5μl of 25mM DTT and 25mM NH_4_HCO_3_ (pH8.5) to 60µl final volume for the reduction step, which was carried out for 15 minutes at 56°C. After cooling on ice to room temperature 3μl of 55mM CAA (chloroacetamide) was added for alkylation where samples were incubated in darkness at room temperature for 15 minutes. This reaction was quenched with 8μl addition of 25mM DTT. Subsequently 15ul of Trypsin/LysC solution [100ng/μl 1:1 Trypsin (Promega):LysC (FujiFilm) mix in 25mM NH_4_HCO_3_] along with 14μl of 25mM NH_4_HCO_3_ (pH8.5) was added to the samples for a final 100µl volume. Digests were carried out overnight at 37°C then subsequently terminated by acidification with 15µl of 2.5% TFA [Trifluoroacetic Acid] to 0.32% final. Samples were desalted and concentrated using Pierce™ C18 SPE pipette tips (100µL volume) per the manufacturer’s protocol. Eluates in 70%:30%:0.1% acetonitrile:water:trifluoroacetic acid were dried to completion then reconstituted in 50µl of 100mM TEAB [Triethylammonium Bicarbonate] for TMT labelling with 10µl [10µg/µl] of reagent [TMTpro 16plex™ Labeling Reagent Set Lot#ZJ391848 from Thermo Scientific in 100% acetonitrile] done at room temperature for 1hr with intermittent gentle vortexing every 15 minutes. Only 10 labels were used for this experiment; channels 126-131N. The labeling reaction was terminated with 2.5µl addition of 5% hydroxylamine [0.2% final] and 15-minute incubation at room temperature. ∼1/12^th^ (5µl, ∼2.4ug) of each individually labelled sample was taken and pooled together for a test pool run to evaluate complexity and mixing ratio. Post test-pool evaluation, master-pool sample was generated where each biological sample was present at the same total protein mass (∼60µg total protein between all samples in ∼155µl total volume), acidified to 1% trifluoroacetic acid, frozen (-80 C) then dried to almost completion in the speed vac. Subsequently resolubilized in 150µl of 5% acetonitrile, 0.3% TFA and 0.2% HFBA for solid-phase extraction using Pierce™ C18 SPE pipette tips (100µl volume) per manufacturer protocol. Enriched samples were solubilized in 35µl of 0.3% TFA, 0.2% HFBA and 3% ACN then desalted using 10µl Pierce® C18 Tips (Thermo Fisher Scientific) according to manufacturer protocol. Eluates in 75%:25%:0.1% acetonitrile:water:TFA acid (vol:vol) were dried to completion in the speed-vac and finally reconstituted in 40ul of 0.1% formic acid and 5% acetonitrile for injection on the instrument.

#### NanoLC-MS/MS

Peptides were analyzed on Orbitrap Fusion™ Lumos™ Tribrid™ platform, where 2µl [∼3µg] total proteome was injected using Dionex UltiMate™3000 RSLCnano delivery system (ThermoFisher Scientific) equipped with an EASY-Spray™ electrospray source (held at constant 50°C). Chromatography of peptides prior to mass spectral analysis was accomplished using capillary emitter column (PepMap® C18, 2µM, 100Å, 500 x 0.075mm, Thermo Fisher Scientific). NanoHPLC system delivered solvents A: 0.1% (v/v) formic acid, and B: 80% (v/v) acetonitrile, 0.1% (v/v) formic acid at 0.30 µL/min to load the peptides at 2% (v/v) B, followed by quick 2 minute gradient to 5% (v/v) B and gradual analytical gradient from 5% (v/v) B to 62.5% (v/v) B over 203 minutes when it concluded with rapid 10 minute ramp to 95% (v/v) B for a 9 minute flash-out. As peptides eluted from the HPLC-column/electrospray source survey MS scans were acquired in the Orbitrap with a resolution of 60,000, max inject time of 50ms and AGC target of 1,000,000 followed by HCD-type MS2 fragmentation into Orbitrap (36% collision energy and 30,000 resolution) with 0.7 m/z isolation window in the quadrupole under ddMSnScan 1 second cycle time mode with peptides detected in the MS1 scan from 400 to 1400 m/z with max inject time of 54ms and AGC target of 125,000; redundancy was limited by dynamic exclusion and MIPS filter mode ON.

#### Data analysis

Raw data was directly imported into Proteome Discoverer 3.1.0.638 where protein identifications and quantitative reporting was generated. Seaquest HT search engine platform with INFERYS rescoring-based workflow for high-resolution MS2 TMT quantification was used to interrogate *Mus musculus* (Mouse) reference proteome database (UP000000589, 04/19/2024 download, 54,874 total entries). Peptide N-terminal and lysine TMTpro labeling plus Cysteine carbamidomethylation were selected as static modifications whereas methionine oxidation was selected as variable modification. Peptide mass tolerances were set at 10ppm for MS1 and 0.03Da for MS2. Peptide and protein identifications were accepted under strict 1% FDR cut offs with high confidence XCorr thresholds of 1.9 for z=2 and 2.3 for z=3. For the total protein quantification processing Reporter Ion Quantifier settings were used on unique and razor peptides, protein grouping was considered for uniqueness. Reporter abundance was based on normalized total peptide amount intensity values with co-isolation threshold filter set at ≤50. ANOVA (individual proteins) hypothesis testing was executed without imputation mode being executed.

#### Proteomics bioinformatics analysis

Raw abundances were normalized to total peptide abundance prior to downstream analysis. Limma 3.64.3 (Ritchie et al. 2015) was used to perform differential expression analysis. All differential expression data can be found in **Supplementary Table 1E**.

#### Functional enrichment analysis

The Gene Set Enrichment Analysis (GSEA) paradigm through its implementation in the R package ‘ClusterProfiler’ v4.16.0, and Bioconductor annotation package ‘org.Mm.eg.db’ v 3.21.0, were used to perform functional enrichment analyses. The DESeq2 t-statistic was used to generate the ranked list of genes. All significant GO terms are reported in **Supplementary Table 1B-D, F, G**.

#### Lipid Extraction

600µL of organic fractions were transferred to 5mL tubes and chilled on ice. Extractions began via the addition of 235µL MeOH (0.01% w/v BHT) which included lipid standards (Avanti Splash II mix, Acylcarnitine NSK-B mix). Samples were then vortexed for 1 minute. This was proceeded by the addition of 750µL MTBE and 1mL of water. Samples were then vortexed for an additional minute, followed by centrifugation at 16100 xg for 5 minutes. 500µL of MTBE were recovered from each sample and dried via speedvac.

#### LC-MSMS Analysis

Dried samples were resuspended in 100µL of IPA. Samples were analyzed using the workflow described in the Agilent application note 5994-3747. In brief, lipids were separated on an Agilent 1290 Infinity II LC coupled to an Agilent 6495C triple quadrupole, using an Agilent Zorbax RRHD Eclipse Plus C18 column (2.1 x 100mm, 1.8µm). Samples were kept in a multisampler at 20°C. The column compartment was held at 45°C. 2µL of extract were injected for each sample. Mobile phase A was comprised of 10mM ammonium formate, 5 µM Agilent deactivator additive in 5:3:2 water:acetonitrile:Isopropanol. Mobile phase B was comprised of 10mM ammonium formate in 1:9:90 water:acetonitrile:isopropanol. The chromatography gradient used the following parameter: starting at 15% B, it increased to 50% at 2.5 minutes, then 57% B at 2.6 minutes, 70% B at 9 minutes, increased to 93% B at 9.1 minutes, increased to 96% B at 11 minutes, increased to 100% B at 11.1 minutes, held at 100% B at 12 minutes, decreased to 15% B at 12.2 minutes, and held at 15% B at 16 minutes for column re-equilibration. The binary pump flow rate was held at 0.4 mL/min.

Regarding source parameters used, this method was operated in polarity-switching mode. The gas temperature was set to 150°C. Drying gas flow rate was held at 17 L/min. Nebulizer gas was held at 20 psi. The sheath gas temperature was held at 200°C with a flow rate of 10 L/min. The capillary voltage was set to 3,500 V and – 3,000V. The nozzle voltage was set to 1,000 V, and -1,500 V. The scan type used was dynamic MRM, utilizing transitions obtained from the Agilent application note 5994-3747EN. Data was collected and analyzed using Agilent MassHunter Suite. Quantification was performed in Agilent MassHunter Workstation and downstream processing was performed using inhouse code using R version 4.1.2 including normalization to internal standards.

#### Lipidomics bioinformatics analysis

Lipids not detected in at least 50% of the samples were removed prior to analysis. Raw abundances were normalized as a percent of all lipids prior to downstream analysis. Limma 3.64.3 (Ritchie et al. 2015) was used to perform differential abundance analysis. All differential abundance data can be found in **Supplementary Data 1H**.

## Statistics

All student’s t-tests were two-tailed. Outliers were identified by ROUT using a threshold of p < 0.05. One-way and two-way ANOVA were conducted assuming a Gaussian distribution and corrected for multiple comparisons using Tukey’s test.

## Contact for Reagent and Resource Sharing

Further information and requests for resources and reagents should be directed to Lead Contact Dr. Rozalyn Anderson.

## Contributions

E.R.M., C.J.M., N.L.A., D.A.B., J.K.N., J.P.C., and K.S.P. generated data and conducted analysis. G.F. and Y.C. maintained the mouse colony. E.R.M., C.J.M., J.A.S., J.F., L.P., and R.M.A. contributed to data interpretation. E.R.M., C.J.M., and R.M.A. wrote the manuscript. All authors contributed to editing the manuscript.

## Acknowledgments

This work was supported by NIH AG057408, AG067330, NIH training fellowship T32DK007665 (ERM), NIH training fellowship T32AG000213 (CJM, DAB, KSP), and the Louis and Elsa Thomsen Distinguished Graduate Fellowship (ERM). Research reported in this publication was supported by the Glenn Foundation and American Federation for Aging Research (A22068 to JAS). The Puglielli laboratory is supported by the NIH (R01NS094154, R01GM148487, and R01AG078794). This study was conducted using resources and facilities at the William S. Middleton Memorial Veterans Hospital, Madison, WI.

## Supplemental Figure Legends

Fig.S1. Pearson rank correlations of all detected lipid species in WT (left) and AT-1^sTg^ neurons (right) following hierarchical clustering.

## Supplemental Table Legends

Table S1. Molecular profiling data. A) Differentially expressed genes (FDR < 5%) between wild type and AT-1^sTg^ neurons by RNA-seq. B) Gene Set Enrichment Analysis Gene Ontology ALL (FDR <5%) for differential regulation between wild type and AT-1^sTg^ neurons by RNA-seq. C) Gene Set Enrichment Analysis Gene Ontology Biological Processes (FDR <5%) for differential regulation between young (10 months) and old (30 months) male mice cortices. D) Gene Set Enrichment Analysis Biological Processes Terms Significant in AT-1 Overexpression and Cortex Aging. E) Differentially abundant proteins (all) between wild type and AT-1^sTg^ neurons by proteomics. F) Gene Set Enrichment Analysis Gene Ontology ALL (p-value<5%) for differential regulation between wild type and AT-1^sTg^ neurons by proteomics. G) Gene Set Enrichment Analysis Terms Significant in RNA-seq and Proteomics. H) Differentially abundant lipids between wild type and AT-1^sTg^ neurons.

## References

Bar S, Wilson KA, Hilsabeck TAU, Alderfer S, Dammer EB, Burton JB, Shah S, Holtz A, Carrera EM, Beck JN, Chen JH, Kauwe G, Seifar F, Shantaraman A, Tracy TE, Seyfried NT, Schilling B, Ellerby LM & Kapahi P (2025) Neuronal glycogen breakdown mitigates tauopathy via pentose-phosphate-pathway-mediated oxidative stress reduction. Nat Metab 7, 1375–1391.

Benavides-Piccione R, Blazquez-Llorca L, Kastanauskaite A, Fernaud-Espinosa I, Tapia-González S & DeFelipe J (2024) Key morphological features of human pyramidal neurons. Cereb Cortex 34, bhae180.

Chang H-C & Guarente L (2014) SIRT1 and other sirtuins in metabolism. Trends Endocrinol Metab 25, 138–145.

Clasquin MF, Melamud E & Rabinowitz JD (2012) LC-MS data processing with MAVEN: a metabolomic analysis and visualization engine. Curr Protoc Bioinformatics Chapter 14, Unit14.11.

Dieterich IA, Lawton AJ, Peng Y, Yu Q, Rhoads TW, Overmyer KA, Cui Y, Armstrong EA, Howell PR, Burhans MS, Li L, Denu JM, Coon JJ, Anderson RM & Puglielli L (2019) Acetyl-CoA flux regulates the proteome and acetyl-proteome to maintain intracellular metabolic crosstalk. Nat Commun 10, 3929.

Duran J, Gruart A, García-Rocha M, Delgado-García JM & Guinovart JJ (2014) Glycogen accumulation underlies neurodegeneration and autophagy impairment in Lafora disease. Hum Mol Genet 23, 3147–3156.

Grimm A & Eckert A (2017) Brain aging and neurodegeneration: from a mitochondrial point of view. Journal of neurochemistry. Available at: 10.1111/jnc.14037.

Hullinger R, Li M, Wang J, Peng Y, Dowell JA, Bomba-Warczak E, Mitchell HA, Burger C, Chapman ER, Denu JM, Li L & Puglielli L (2016) Increased expression of AT-1/SLC33A1 causes an autistic-like phenotype in mice by affecting dendritic branching and spine formation. Journal of Experimental Medicine 213, 1267–1284.

Jankowska-Kulawy A, Klimaszewska-Łata J, Gul-Hinc S, Ronowska A & Szutowicz A (2022) Metabolic and Cellular Compartments of Acetyl-CoA in the Healthy and Diseased Brain. Int J Mol Sci 23, 10073.

Jonas MC, Pehar M & Puglielli L (2010) AT-1 is the ER membrane acetyl-CoA transporter and is essential for cell viability. Journal of Cell Science 123, 3378–3388.

Khaliulin I, Hamoudi W & Amal H (2025) The multifaceted role of mitochondria in autism spectrum disorder. Mol Psychiatry 30, 629–650.

Kirchner JH, Euler L & Gjorgjieva J (2023) Dendritic growth and synaptic organization from activity-independent cues and local activity-dependent plasticity. eLife 12. Available at: https://elifesciences.org/reviewed-preprints/87527 [Accessed September 2, 2024].

Ko MH & Puglielli L (2009) Two endoplasmic reticulum (ER)/ER Golgi intermediate compartment-based lysine acetyltransferases post-translationally regulate BACE1 levels. J Biol Chem 284, 2482–2492.

Lee A, Hirabayashi Y, Kwon S-K, Lewis TL & Polleux F (2018) Emerging roles of mitochondria in synaptic transmission and neurodegeneration. Current Opinion in Physiology 3, 82–93.

Liao Y, Smyth GK & Shi W (2019) The R package Rsubread is easier, faster, cheaper and better for alignment and quantification of RNA sequencing reads. Nucleic Acids Research 47, e47.

Lilja J & Ivaska J (2018) Integrin activity in neuronal connectivity. J Cell Sci 131, jcs212803.

Maier T, Guell M & Serrano L (2009) Correlation of mRNA and protein in complex biological samples. FEBS letters 583, 3966–73.

McGregor ER, Lasky DJ, Rippentrop OJ, Clark JP, Wright S, Jones MV & Anderson RM (2025) Reversal of neuronal tau pathology via adiponectin receptor activation. Commun Biol 8, 8.

Melamud E, Vastag L & Rabinowitz JD (2010) Metabolomic analysis and visualization engine for LC-MS data. Anal Chem 82, 9818–9826.

Miller KN, Clark JP & Anderson RM (2019) Mitochondrial regulator PGC-1a-Modulating the modulator. Curr Opin Endocr Metab Res 5, 37–44.

Miller KN, Clark JP, Martin SA, Howell PR, Burhans MS, Haws SA, Johnson NB, Rhoads TW, Pavelec DM, Eliceiri KW, Roopra AS, Ntambi JM, Denu JM, Parks BW & Anderson RM (2019) PGC-1a integrates a metabolism and growth network linked to caloric restriction. Aging Cell 18. Available at: https://onlinelibrary.wiley.com/doi/10.1111/acel.12999 [Accessed July 16, 2022].

Monzel AS, Enríquez JA & Picard M (2023) Multifaceted mitochondria: moving mitochondrial science beyond function and dysfunction. Nat Metab 5, 546–562.

Peng Y, Li M, Clarkson BD, Pehar M, Lao PJ, Hillmer AT, Barnhart TE, Christian BT, Mitchell HA, Bendlin BB, Sandor M & Puglielli L (2014) Deficient import of acetyl-CoA into the ER lumen causes neurodegeneration and propensity to infections, inflammation, and cancer. J Neurosci 34, 6772–89.

Peng Y, Shapiro SL, Banduseela VC, Dieterich IA, Hewitt KJ, Bresnick EH, Kong G, Zhang J, Schueler KL, Keller MP, Attie AD, Hacker TA, Sullivan R, Kielar-Grevstad E, Arriola Apelo SI, Lamming DW, Anderson RM & Puglielli L (2018) Increased transport of acetyl-CoA into the endoplasmic reticulum causes a progeria-like phenotype. Aging Cell 17, e12820.

Picard M, Shirihai OS, Gentil BJ & Burelle Y (2013) Mitochondrial morphology transitions and functions: implications for retrograde signaling? Am J Physiol Regul Integr Comp Physiol 304, R393–R406.

Pietrocola F, Galluzzi L, Bravo-San Pedro JM, Madeo F & Kroemer G (2015) Acetyl coenzyme A: a central metabolite and second messenger. Cell Metab 21, 805–821.

Rai A, Singh PK, Singh V, Kumar V, Mishra R, Thakur AK, Mahadevan A, Shankar SK, Jana NR & Ganesh S (2018) Glycogen synthase protects neurons from cytotoxicity of mutant huntingtin by enhancing the autophagy flux. Cell Death Dis 9, 201.

Ritchie ME, Phipson B, Wu D, Hu Y, Law CW, Shi W & Smyth GK (2015) limma powers differential expression analyses for RNA-sequencing and microarray studies. Nucleic Acids Res 43, e47.

Saez I, Duran J, Sinadinos C, Beltran A, Yanes O, Tevy MF, Martínez-Pons C, Milán M & Guinovart JJ (2014) Neurons Have an Active Glycogen Metabolism that Contributes to Tolerance to Hypoxia. J Cereb Blood Flow Metab 34, 945–955.

Schaefer PM, Kalinina S, Rueck A, von Arnim CAF & von Einem B (2019) NADH Autofluorescence-A Marker on its Way to Boost Bioenergetic Research: NADH Autofluorescence. Cytometry 95, 34–46.

Sekiya M, Kainoh K, Sugasawa T, Yoshino R, Hirokawa T, Tokiwa H, Nakano S, Nagatoishi S, Tsumoto K, Takeuchi Y, Miyamoto T, Matsuzaka T & Shimano H (2021) The transcriptional corepressor CtBP2 serves as a metabolite sensor orchestrating hepatic glucose and lipid homeostasis. Nat Commun 12, 6315.

Shi L & Tu BP (2015) Acetyl-CoA and the regulation of metabolism: mechanisms and consequences. Curr Opin Cell Biol 33, 125–131.

Souder DC, McGregor ER, Clark JP, Rhoads TW, Porter TJ, Eliceiri KW, Moore DL, Puglielli L & Anderson RM (2025) Neuron-specific isoform of PGC-1α regulates neuronal metabolism and brain aging. Nat Commun 16, 2053.

Soyal SM, Bonova P, Kwik M, Zara G, Auer S, Scharler C, Strunk D, Nofziger C, Paulmichl M & Patsch W (2020) The Expression of CNS-Specific PPARGC1A Transcripts Is Regulated by Hypoxia and a Variable GT Repeat Polymorphism. Mol Neurobiol 57, 752–764.

Thio SSC, Bonventre JV & Hsu SI-H (2004) The CtBP2 co-repressor is regulated by NADH-dependent dimerization and possesses a novel N-terminal repression domain. Nucleic Acids Res 32, 1836–1847.

Turnbull J, DePaoli-Roach AA, Zhao X, Cortez MA, Pencea N, Tiberia E, Piliguian M, Roach PJ, Wang P, Ackerley CA & Minassian BA (2011) PTG depletion removes Lafora bodies and rescues the fatal epilepsy of Lafora disease. PLoS Genet 7, e1002037.

Wang D, Eraslan B, Wieland T, Hallström B, Hopf T, Zolg DP, Zecha J, Asplund A, Li L-H, Meng C, Frejno M, Schmidt T, Schnatbaum K, Wilhelm M, Ponten F, Uhlen M, Gagneur J, Hahne H & Kuster B (2019) A deep proteome and transcriptome abundance atlas of 29 healthy human tissues. Mol Syst Biol 15, e8503.

Xu X, Pang Y & Fan X (2025) Mitochondria in oxidative stress, inflammation and aging: from mechanisms to therapeutic advances. Signal Transduct Target Ther 10, 190.

Zhao K, Hong H, Zhao L, Huang S, Gao Y, Metwally E, Jiang Y, Sigrist SJ & Zhang YQ (2020) Postsynaptic cAMP signalling regulates the antagonistic balance of Drosophila glutamate receptor subtypes. Development 147, dev191874.

Zhu Y, Fan Z, Wang R, Xie R, Guo H, Zhang M, Guo B, Sun T, Zhang H, Zhuo L, Li Y & Wu S (2020) Single-Cell Analysis for Glycogen Localization and Metabolism in Cultured Astrocytes. Cell Mol Neurobiol 40, 801–812.

