## Supplementary figures and images for "Acetyl-CoA availability regulates neuronal metabolism, growth, and synaptic activity"

### Supplemental Figure 1

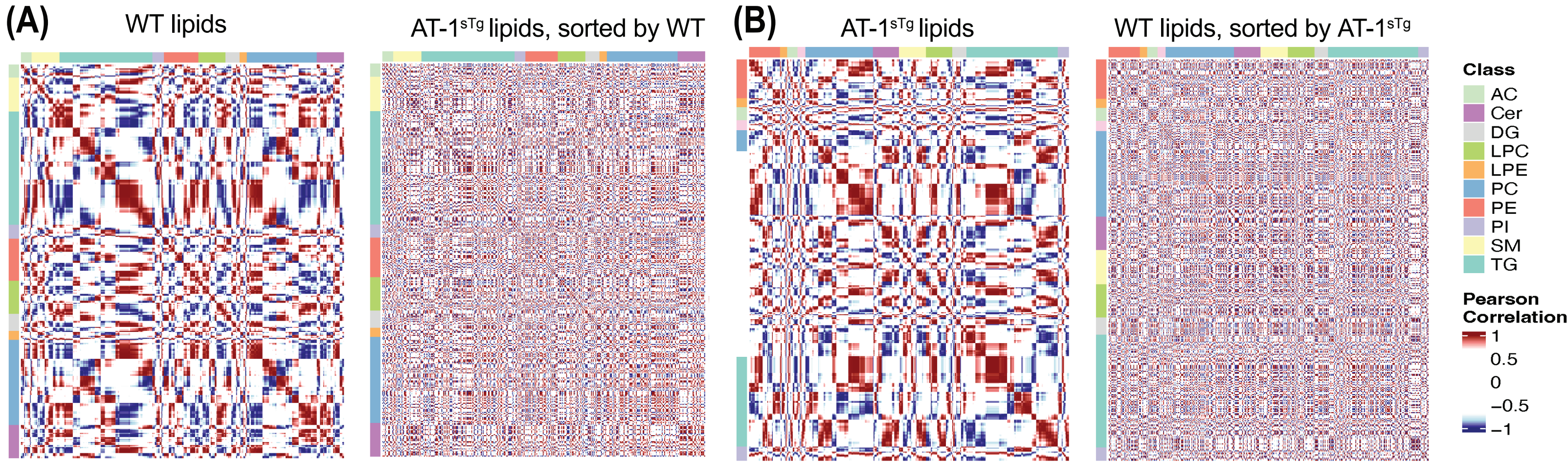
